# Temporal attention improves perception without enhancement by temporal competition

**DOI:** 10.64898/2026.02.11.705419

**Authors:** Karen J. Tian, Jennifer A. Motzer, Rachel N. Denison

## Abstract

When successive stimuli occur close enough together in time, their perception can be impaired. Such impairments indicate temporal competition between successive stimuli for processing resources. Voluntary temporal attention can bias processing resources in favor of a behaviorally relevant moment, improving perception at the attended time at the expense of impairments at unattended times. However, it is unclear whether these perceptual tradeoffs across time arise because voluntary temporal attention selects among actively competing stimulus representations, such as within visual working memory, or if instead, temporal attention facilitates stimulus processing prior to a competitive stage. Here we used a temporal cueing task with up to two grating targets in succession to test whether and how the effects of temporal attention depend on temporal competition. Participants reported whether at a probed time the target was present or absent, and if present, performed a difficult orientation discrimination judgment. To isolate the effects of voluntary temporal attention from those of temporal expectation, we manipulated the task-relevance of the two possible target times while controlling stimulus timing predictability. Temporal competition was manipulated via stimulus presence or absence at the unprobed time. We found that voluntary temporal attention improved perceptual discriminability even in the absence of temporal competition, when only one stimulus appeared during the trial. Critically, the magnitude of attentional enhancement was comparable with and without temporal competition. These results suggest that voluntary temporal attention enhances perception by facilitating processing prior to a competitive stage, rather than by resolving conflicts between actively competing stimulus representations.

**Graphical abstract:** 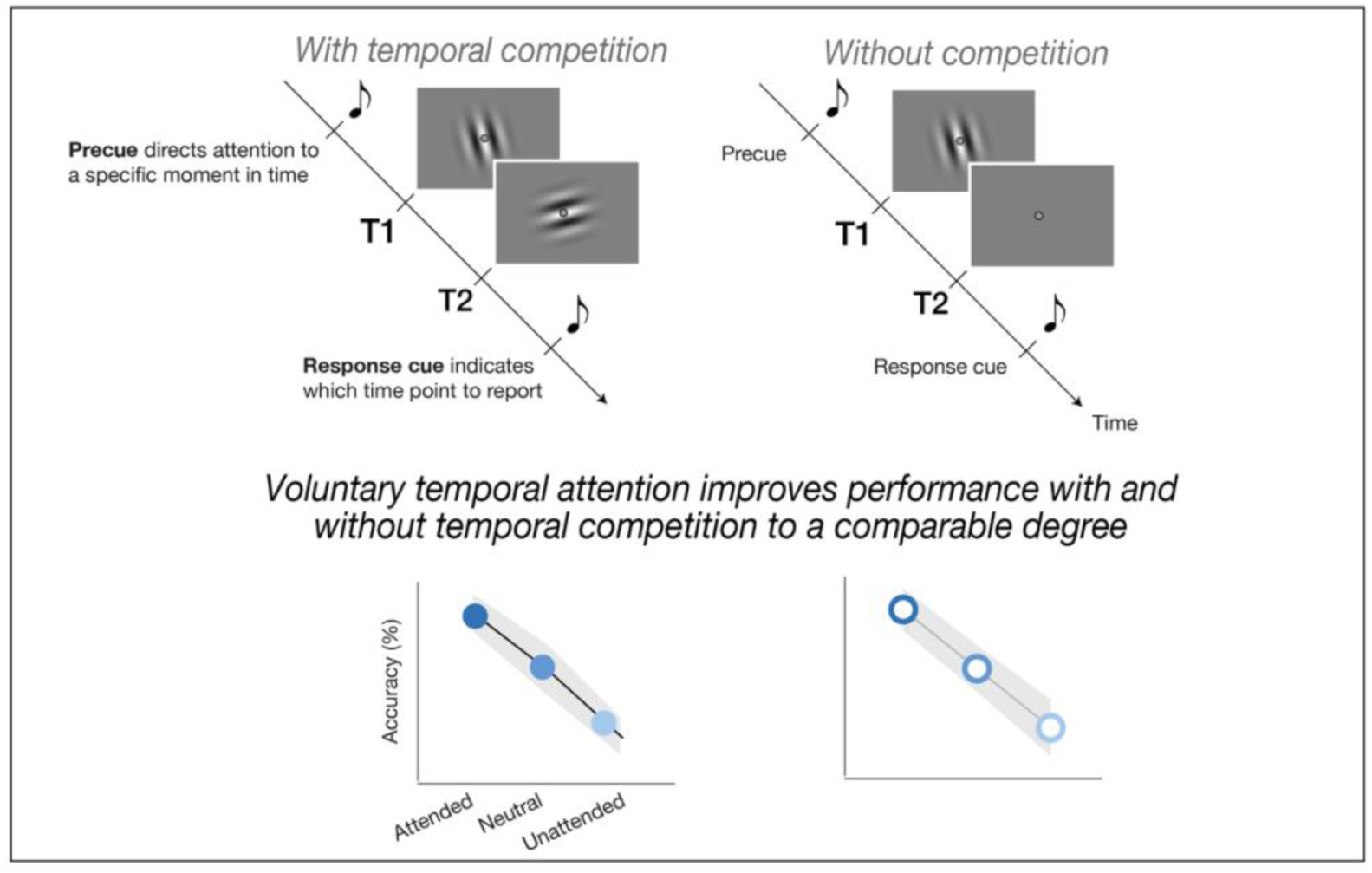

## Introduction

Visual input is continuous, yet visual processing is limited. Perception can be impaired when successive stimuli appear close together in time (Breitmeyer & Öğmen, 2006; Epstein et al., 2025; Hochmitz et al., 2024; Sahar & Yeshurun, 2023; Shapiro & Raymond, 2025), demonstrating temporal competition for processing resources. Such processing constraints can be managed by attention. Voluntary temporal attention is the deliberate prioritization of sensory information at a specific moment in time known in advance to be relevant to one’s behavioral goals (Denison et al., 2017; Denison, 2024; Nobre & van Ede, 2018, 2023; Seibold et al., 2023).

When stimuli compete in time, voluntary temporal attention produces perceptual tradeoffs: improved perception at the attended time point, at the expense of impaired perception before and after (Denison et al., 2017; Denison, 2024; Nobre & van Ede, 2018, 2023). These tradeoffs are most pronounced when successive stimuli are separated by 200 to 450 ms (Denison et al., 2017, 2024; Duyar et al., 2024; Fernández et al., 2019; Fu et al., 2026; Huang et al., 2025; Jing et al., 2023; Palmieri et al., 2023; Zhu et al., 2024) and disappear at longer intervals of 800 ms (Denison et al., 2021). A similar temporal profile characterizes perceptual interference between successive stimuli in the absence of an attentional manipulation (Hochmitz et al., 2026; Sahar & Yeshurun, 2023; Tkacz-Domb & Yeshurun, 2017, 2021). Taken together, these findings could indicate that the effects of temporal attention critically depend on the presence of temporally competing stimuli.

On the other hand, a single stimulus presented at a more versus less expected time is processed more quickly (Capizzi et al., 2023; Correa et al., 2004; Correa, Lupiáñez, & Tudela, 2006; Coull et al., 2000; Coull, 2004; Coull & Nobre, 1998; Griffin et al., 2001; Miniussi et al., 1999; Naccache et al., 2001; Nobre, 2001) and accurately (Chauvin et al., 2016; Rohenkohl et al., 2011). Such results might be taken to indicate that temporal attention does not depend on temporal competition, but no study has directly compared the magnitudes of attentional effects with and without temporal competition. Moreover, single-stimulus protocols have not isolated temporal attention—associated with the behavioral relevance of a certain moment—from temporal expectation—associated with the predictability of stimulus timing (Denison, 2024). Whether temporal attention requires competition when expectation is controlled has not been investigated.

Comparing the effects of voluntary temporal attention on perception with and without temporal competition can reveal the stage of processing at which temporal attention facilitates perception. Temporal attention could facilitate processing before an active competitive stage, for example, by transiently modulating sensory gain at the attended moment. Alternatively, temporal attention could facilitate perception during a competitive stage, biasing selection among actively competing representations in the ventral stream or in working memory. A critical role of competition in eliciting attentional effects has been emphasized in the spatial domain: the influential biased competition framework (Desimone, 1998; Desimone & Duncan, 1995) proposes that attention resolves competition among simultaneous stimuli by biasing representational resources in favor of the behaviorally relevant stimulus. By analogy, temporal attention may mediate biased competition across time.

These mechanistic accounts predict effects of temporal attention on perception that are indistinguishable when temporal competition is present but diverge when competition is absent. If temporal attention facilitates sensory processing before a competitive stage, then it should improve the perception of a single stimulus to the same degree when competition is present or absent **(Figure 1a)**. If instead temporal attention acts both before and during an active competitive stage, then it should improve the perception of a single stimulus, but to a lesser degree than when competition is present **(Figure 1b)**. Finally, if temporal attention exclusively selects among actively competing representations, then it should not affect the perception of a single stimulus, which faces no competition for attention to resolve **(Figure 1c)**.

**Figure 1.**
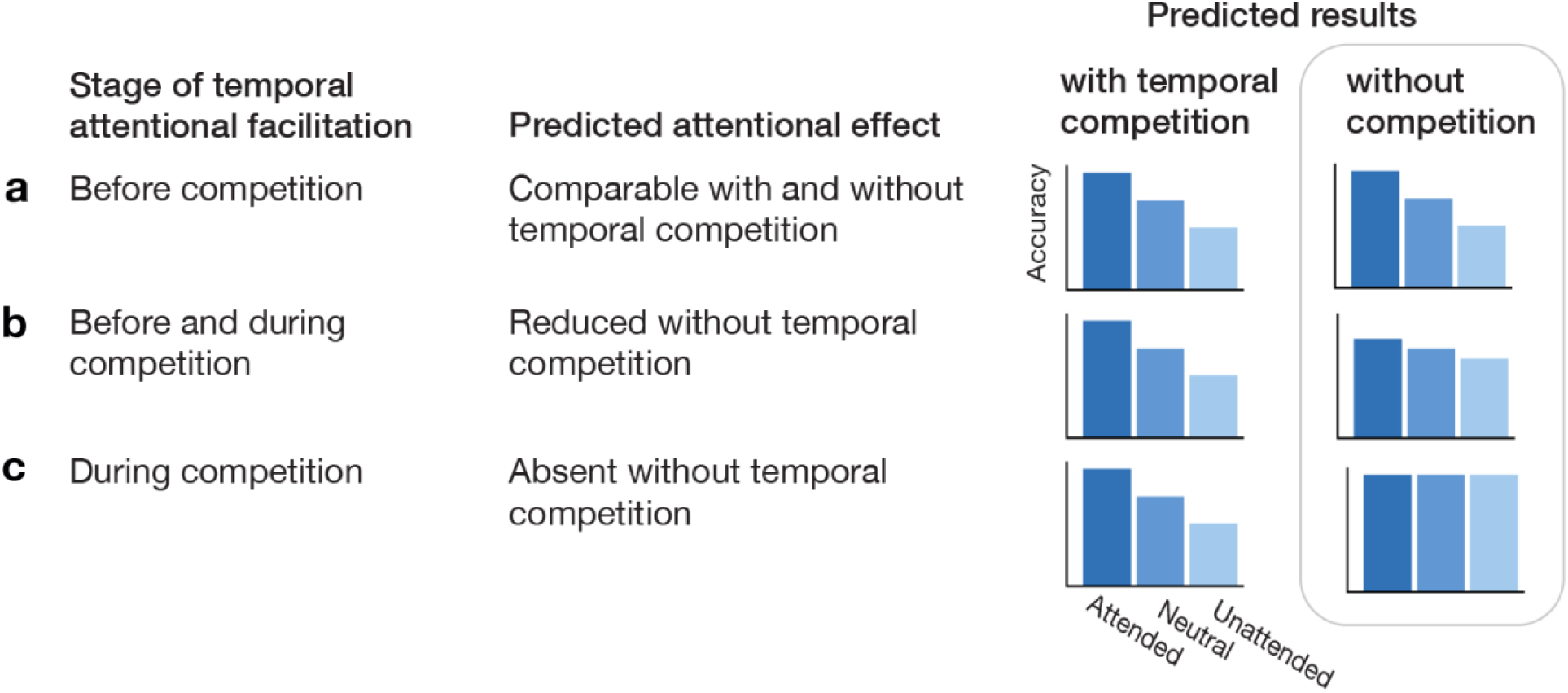
Hypothesized mechanisms of voluntary temporal attention and predicted behavioral outcomes. Voluntary temporal attention could facilitate perception **a)** before an active competitive stage, **b)** before and during a competitive stage, or **c)** only during a competitive stage. These mechanistic accounts predict behavioral effects of temporal attention that are indistinguishable when temporal competition is present (left column of bars) but diverge when temporal competition is absent (right column of bars). Depending on the underlying mechanism, the effects of voluntary temporal attention for a single stimulus without temporal competition should (a) persist and be comparable in magnitude to when temporal competition is present, (b) persist but be reduced in magnitude, or (c) be absent.

To arbitrate among these mechanistic accounts, here we designed a task that independently manipulated voluntary temporal attention and temporal competition while controlling temporal expectation. We found that voluntary temporal attention improved perceptual discriminability to a similar degree with and without temporal competition. Thus, the perceptual enhancements of voluntary temporal attention neither required nor were amplified by temporal competition, supporting an account in which temporal attention facilitates sensory processing before an active competitive stage.

## Results

We independently manipulated temporal attention and temporal competition during an orientation judgment task. On each trial, gratings could appear at two successive, predictable time points, T1 and T2 **(Figure 2)**, separated by 250 ms. These time points were marked by auditory clicks and fixation dims, minimizing temporal uncertainty. A subsequent response cue indicated which time point to report using a three-way judgment: grating present and clockwise (CW), present and counterclockwise (CCW), or absent. We refer to the time point probed by the response cue as the “target” and the other time point as the “non-target”. Temporal attention was manipulated via a precue that directed participants to attend to T1 or T2 with 75% validity, or was uninformative on neutral precue trials. Temporal competition was manipulated via the presence or absence of a grating at the non-target time point. Gratings were equally likely to be present or absent at each time point, independently for T1 and T2.

**Figure 2.**
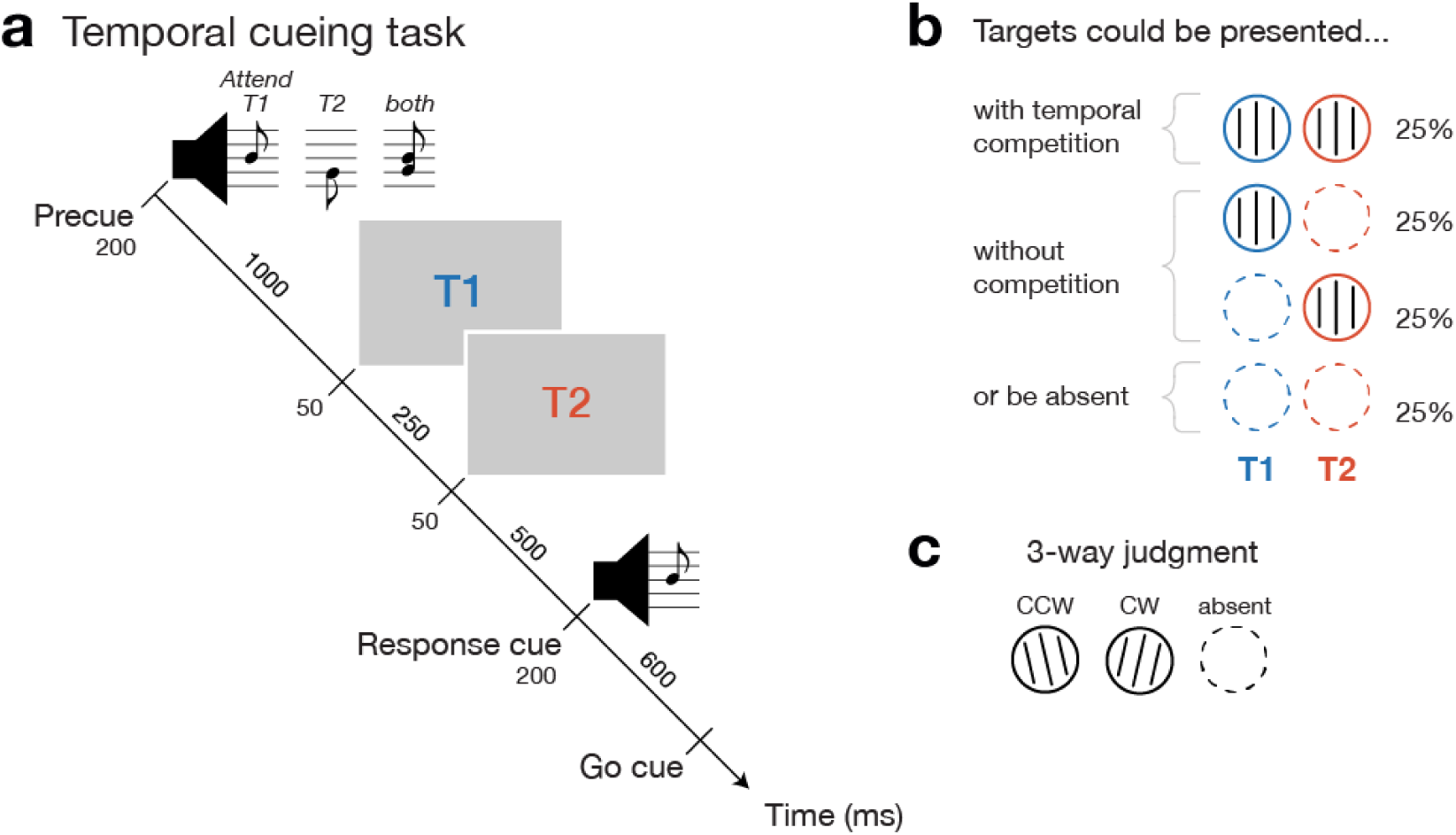
Temporal cueing task. **a)** Trial timeline. Each trial contained two time points (T1, T2) at which gratings could appear. An auditory precue directed participants’ attention to a single time point (as indicated by a single tone) or distributed attention across time points (tones together, neutral attention condition). A response cue indicated which single time point to report, matching informative precues with 75% validity. **b)** Possible stimulus sequences. On a given trial, gratings could appear at both time points (generating temporal competition), at only one time point (without competition), or at neither time point. Trial percentages are shown for each sequence type. **c)** Participants made a three-way perceptual judgment, reporting whether or not they saw a grating at the response-cued time point, and if so, its tilt (CCW=counterclockwise, CW=clockwise). Gratings were slightly tilted from either the vertical or horizontal axis, with independent presence, axis, and tilt for T1 and T2. Grating schematics are shown vertically oriented for illustration only.

### Voluntary temporal attention generated perceptual tradeoffs

To confirm the efficacy of the attentional manipulation, we first assessed accuracy on the three-way perceptual judgment (“CCW, CW, or absent?”) as a function of the precue condition, irrespective of the non-target. When the target was present **(Figure 3a)**, accuracy across target time points was highest for valid, intermediate for neutral, and lowest for invalid conditions (main effect of validity: *F*(2,34)=28.74, *p*<0.001, *η*_G_^2^=0.15), demonstrating temporal attentional benefits and costs on perception relative to the neutral condition (Denison et al., 2017, 2021, 2024; Fernández et al., 2019; Jing et al., 2023; Zhu et al., 2024). Because of the directional nature of time, we also examined each target separately; unless otherwise noted, target-specific statistics are reported in the Supplementary Material. Temporal attention improved performance for each target individually (Supplementary Table 1) and more so for T1 (validity x target: *F*(2,34)=4.31, *p*=0.022, *η*_G_^2^=0.03), possibly reflecting the overall higher accuracy for T2 (main effect of target: *F*(1,17)=69.12, *p*<0.001, *η*_G_^2^=0.23). Temporal attention also decreased reaction times (**Figure 3c)** (*F*(2,34)=18.20, *p*<0.001, *η*_G_^2^=0.07; Supplementary Table 2), ruling out a speed-accuracy trade-off in driving perceptual improvements.

**Figure 3.**
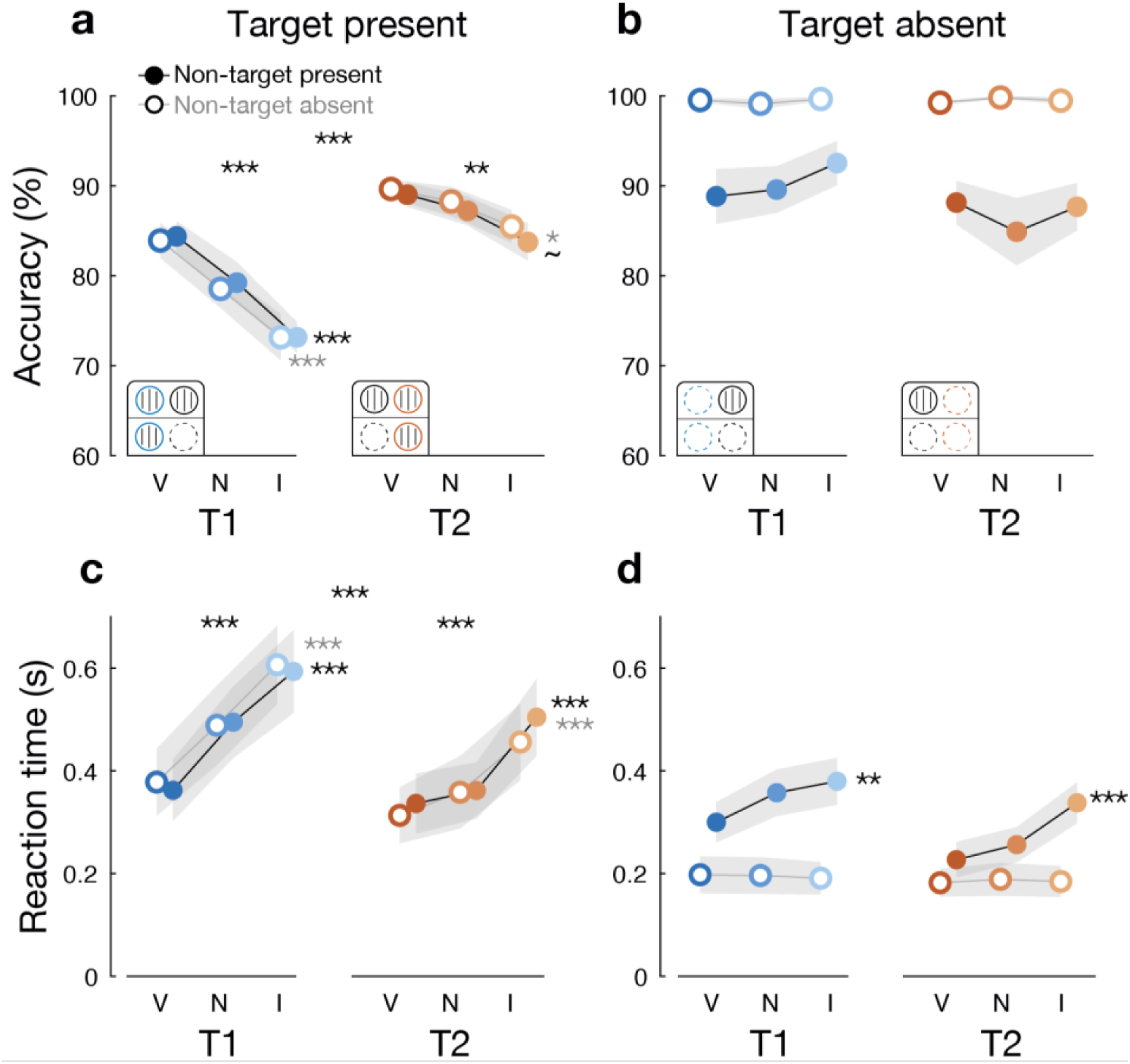
Temporal attentional improvements to perceptual accuracy and reaction time. **a,b)** Accuracy and **c,d)** reaction time on the three-way perceptual judgment as a function of target presence and non-target presence. Grating icons show possible trial sequences for each condition; e.g., the target present T1 condition at left includes both T1 present / T2 absent trials and T1 present / T2 present trials. Data (n=18) are shown as group means ±1 SEM. Stars indicate significance levels for effects of validity only in cases where there was no interaction with temporal competition. V=valid, N=neutral, I=invalid; ∼*p*<0.1, \**p*<0.05, \*\**p*<0.01, \*\*\**p*<0.001.

When the target was absent **(Figure 3b)**, accuracy was overall high (mean=94.04%, SD=6.13%) and was not modulated by temporal attention (no effect of validity: *F*(2,34)=2.33, *p*=0.113, *η*_G_^2^<0.01, Supplementary Table 1). Temporal attention thus improved the challenging tilt discrimination of a present target but not the near-ceiling accuracy in labeling an absent target (validity x target presence: *F*(2,34)=35.23, *p*<0.001, *η*_G_^2^=0.05). Temporal attention speeded reaction times on target-absent trials overall **(Figure 3d)** (main effect of validity: *F*(2,34)=5.42, *p*=0.009, *η*_G_^2^=0.02; Supplementary Table 2), likely reflecting the time it takes for participants to update their response plan following an invalid precue.

### Comparable temporal attentional tradeoffs with and without temporal competition

To assess whether the magnitude of temporal attentional effects depends on temporal competition, we examined accuracy as a function of the non-target. When the target was present (**Figure 3a)**, temporal attention improved accuracy both when the non-target was present (main effect of validity: *F*(2,34)=10.40, *p*<0.001, *η*_G_^2^=0.17; Supplementary Table 1) and when it was absent (*F*(2,34)=14.98, *p*<0.001, *η*_G_^2^=0.13).

Critically, there was no significant interaction between validity and non-target presence (*F*(2,34)=0.06, *p*=0.941). Because one hypothesis predicted no interaction of temporal attention and temporal competition **(Figure 1a)**, we conducted planned Bayesian statistics to distinguish insufficient statistical power from evidence in support of the null. This analysis yielded a BF _10_ of 0.084, indicating strong evidence in favor of the null (Lee & Wagenmakers, 2014). Therefore, temporal attention generates perceptual tradeoffs in the absence of temporal competition, with effects comparable in magnitude to when competition was present, supporting a mechanism in which temporal attention facilitates perception before an active competitive stage.

When the target was absent **(Figure 3b)**, accuracy was impaired by the presence of a non-target (*F*(1,17)=21.31, *p*<0.001, *η*_G_^2^=0.30) but unaffected by temporal attention, whether the non-target was present or absent (all *F*(2,34)<2.35, *p*>0.110). Reaction times were slower when the non-target was present **(Figure 3d)** (*F*(1,17)=35.70, *p*<0.001, *η*_G_^2^=0.14) and faster with attention (*F*(2,34)=5.42, *p*=0.009, *η*_G_^2^=0.02), and this attentional speeding depended on the non-target (validity x non-target presence: F(2,34)=10.62, p<0.001, *η*_G_^2^=0.02). On trials in which both stimuli were absent, we found no effect of attention on accuracy or reaction time (Supplementary Tables 2 and 3), likely reflecting an easy decision when no gratings were presented.

Because target detection and fine tilt discrimination are combined in the three-way judgement, we next used signal detection theory (SDT) (Green & Swets, 1966) to assess whether temporal attention and temporal competition exert separable effects on detection and discrimination.

### Temporal competition impaired discrimination sensitivity but did not interact with temporal attention

We assessed the effects of voluntary temporal attention and temporal competition on tilt discrimination, which was necessarily restricted to trials in which the target was present and a tilt was reported. Because the number of such trials differed across attention conditions, and unequal trial counts could differentially affect SDT estimates, we applied a subsampling procedure to all SDT analyses to control trial counts (see Methods, SDT analysis).

Discrimination sensitivity was improved by temporal attention **(Figure 4)** (main effect of validity: *F*(2,34)=19.44, *p*<0.001, *η*_G_^2^=0.10; Supplementary Table 3; Supplementary Note 1) and was impaired by temporal competition (main effect of non-target presence: *F*(1,17)=22.61, *p*<0.001, *η*_G_^2^=0.13). Critically, temporal attention improved discrimination sensitivity both when the non-target was present (*F*(2,34)=11.53, *p*<0.001, *η*_G_^2^=0.18) and when it was absent (*F*(2,34)=4.36, *p*=0.021, *η*_G_^2^=0.04), with no evidence that the magnitude of attentional effects differed with the non-target (no validity x non-target presence interaction: *F*(2,34)=2.45, *p*=0.101, *η*_G_^2^=0.02, BF_10_=0.908). Thus temporal attention altered the fidelity of perceptual representations with and without temporal competition.

**Figure 4.**
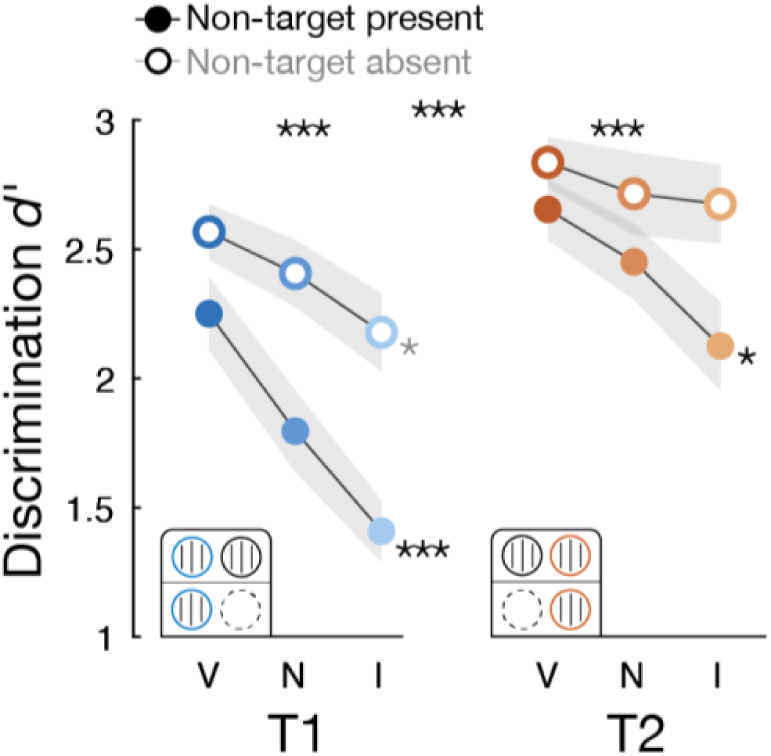
Temporal attention improved discrimination sensitivity. Discrimination sensitivity (*d′*) was computed from trials in which the target was present, and a tilt response was supplied. **a)** Temporal attention enhanced discrimination sensitivity and temporal competition reduced it, with no significant interaction. Data (n = 18) are shown as group means ±1 SEM. Stars indicate effects of validity. V=valid, N=neutral, I=invalid; \**p*<0.05, \*\*\**p*<0.001.

### Temporal attention improved detection sensitivity when the non-target was absent

To assess whether temporal attention and competition also influence stimulus detection, we calculated detection sensitivity—the ability to distinguish target-present vs. -absent trials— labeling both “CW” and “CCW” tilt responses as target-present reports. Because gratings were full contrast, detection errors likely reflected temporal order confusion, such as reporting T1 as present when only T2 was present.

Detection sensitivity was overall high **(Figure 5a)** (mean *d’*=3.63, SD=0.54), as expected for suprathreshold gratings, and higher for T2 than T1 (main effect of target: *F*(1,17)=5.78, *p*=0.028, *η*_G_^2^=0.02). Detection sensitivity was impaired by the non-target (main effect of non-target presence: *F*(1,17)=6.12, *p*=0.024, *η*_G_^2^=0.05), which was specific to T2 (target x non-target presence: *F*(1,17)=16.50, *p*<0.001, *η*_G_^2^=0.09).

**Figure 5.**
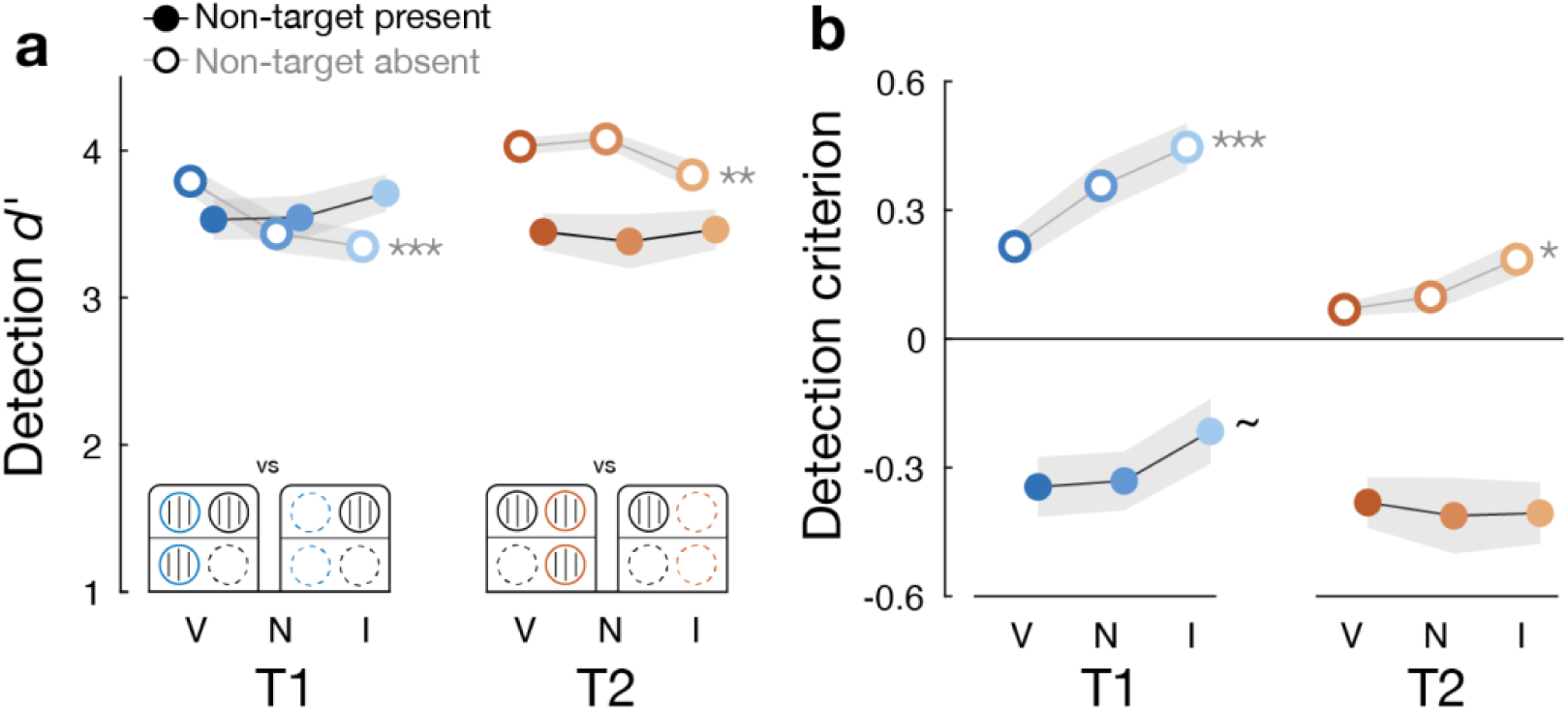
Temporal attentional effects on detection sensitivity. Detection sensitivity (d′) and criterion for distinguishing whether the target was present or absent. a) Detection sensitivity was impaired by the presence of a non-target, and enhanced by temporal attention only when the non-target was absent. b) Detection criterion was strongly influenced by the non-target, with more liberal (i.e., lower) values when the non-target was present. Data (n=18) are shown as group means ±1 SEM. Stars indicate significance levels for effects of validity. V=valid, N=neutral, I=invalid; ∼p<0.1, *p<0.05, **p<0.01, ***p<0.001.

In contrast to its effects on discrimination sensitivity, the effects of temporal attention on detection sensitivity depended on the non-target (validity x non-target presence: *F*(2,34)=11.10, *p*<0.001, *η*_G_^2^=0.03; Supplementary Table 4). Interestingly, temporal attention improved detection sensitivity only when the non-target was absent (main effect of validity: *F*(2,34)=10.48, *p*<0.001, *η*_G_^2^=0.10) and not when it was present (*F*(2,34)=1.35, *p=*0.273, *η*_G_^2^<0.01). This finding could indicate that, when only one stimulus is presented, temporal attention improves the perception of stimulus timing.

Detection criterion—the propensity to report seeing a target—was strongly modulated by the non-target (main effect of non-target presence: *F*(1,17)=66.39, *p*<0.001, *η*_G_^2^=0.58). Criterion was liberal (negative values) when a non-target was present, but conservative (positive values) when the non-target was absent **(Figure 5b)**. Participants’ detection reports were therefore biased toward consistency with the non-target, and this tendency was stronger for T1 (target x non-target presence: *F*(1,17)=4.54, *p*=0.048, *η*_G_^2^=0.02), despite the independence of the target and non-target. This bias may reflect temporal order errors or temporal integration, at perceptual or decisional levels.

Temporal attention induced a more liberal detection criterion overall (i.e., lower values) (main effect of validity: *F*(2,34)=12.52, *p*<0.001, *η*_G_^2^=0.04; Supplementary Table 4), indicating participants had a greater propensity to report a target as present when it was attended. This shift was pronounced when the non-target was absent (*F*(2,34)=13.66, *p*<0.001, *η*_G_^2^=0.14); conversely, when the non-target was present, attention had no significant effect on criterion (*F*(2,34)=1.19, *p*=0.316). Thus temporal attention made the detection criterion more liberal, particularly when the non-target was absent.

## Discussion

We investigated whether the effects of voluntary temporal attention on perception depend on temporal competition. Temporal attention generated perceptual tradeoffs—enhanced perceptual discriminability at the attended time point and impairments at the unattended time point relative to a neutral condition—that were comparable in magnitude for a stimulus presented alone vs. with a temporally proximal competitor. These findings are inconsistent with models in which attention acts solely to resolve active competition, and instead support a mechanism in which attention facilitates processing at a task-relevant moment before a competitive stage.

The perception of a stimulus can be influenced by temporal context extending hundreds of milliseconds into the past and future (Akyürek, 2025; Chapman & Denison, 2025; Epstein et al., 2025; Fleming & Michel, 2025; Herzog et al., 2020; Sergent et al., 2013). In temporal crowding, perception is impaired when other stimuli appear within a temporal window of 150-450 ms (Hochmitz et al., 2026; Sahar & Yeshurun, 2023; Tkacz-Domb & Yeshurun, 2017, 2021). In the attentional blink, perception is impaired 200-500 ms following an initial target that reactively elicits attention (Broadbent & Broadbent, 1987; Raymond et al., 1992; Shapiro & Raymond, 2025). When attention is instead strategically deployed in advance, as in voluntary temporal attention, perceptual effects are most pronounced for stimuli separated by 200-450 ms (Denison et al., 2021). These phenomena, identified using distinct experimental tasks, operate on a remarkably similar timescale, suggesting they may be different expressions of a common processing bottleneck.

In the biased competition framework, developed for static images, spatial attention resolves competition between simultaneous stimuli for limited representational resources (Desimone, 1998; Desimone & Duncan, 1995). Motivating this view, when multiple stimuli fall within the spatial receptive field of a visual cortical neuron, attention biases the neuron’s response towards what it would be if only the attended stimulus were present (Chelazzi et al., 1993, 2001; Luck et al., 1997; Moran & Desimone, 1985; Reynolds et al., 1999, 2000; Reynolds & Chelazzi, 2004); for a single stimulus, attentional modulations are weaker or absent (Chelazzi et al., 2001; Ito & Gilbert, 1999; Luck et al., 1997; Moran & Desimone, 1985; Motter, 1993; Posner & Gilbert, 1999; Reynolds et al., 1999, 2000). Behaviorally, spatial attention still improves the perception of a single stimulus (Cameron et al., 2002; Carrasco et al., 2000, 2002; Jigo & Carrasco, 2020; Ling & Carrasco, 2006), but typically to a lesser degree than when multiple stimuli compete across space (Carrasco & McElree, 2001; Foley & Schwarz, 1998; Giordano et al., 2009; Palmer, 1994). These patterns are consistent with the normalization model of attention (Reynolds & Heeger, 2009), in which attention shifts the balance of excitation and suppression between simultaneous stimuli, thereby exerting greater modulatory influence when competitive interactions are stronger.

Extending biased competition to dynamic sequences, we can consider at least two possible neural sites of temporal competition: 1) ventral stream regions with long temporal integration windows (Chaudhuri et al., 2015; Hasson et al., 2008; Honey et al., 2012; Murray et al., 2014), and 2) visual working memory, where sequential items are concurrently maintained (Shalev & van Ede, 2021). Temporal attention could mediate competitive interactions by biasing relevant stimulus representations at one or both of these sites. Instead, we found that voluntary temporal attention improved perceptual discriminability similarly with and without a temporally competing stimulus, suggesting that temporal attentional modulations occur upstream of these competitive sites.

Physiological evidence indicates that voluntary temporal attention facilitates processing in anticipation of the first stimulus (Denison et al., 2019, 2024; Duyar & Carrasco, 2025; Palmieri et al., 2023) and before a second stimulus (Denison et al., 2024; Zhu et al., 2024). These studies suggest neural mechanisms by which temporal attention could impact processing before competition, including periodic attentional modulations (Denison et al., 2024) and selective routing of visual signals (Zhu et al., 2024, 2026). Some studies that combine temporal attention and expectation have also identified early modulations (Anderson & Sheinberg, 2008; Correa, Lupiáñez, Madrid, et al., 2006; Griffin et al., 2002; Lima et al., 2011; Miniussi et al., 1999). However, these physiological studies could not determine the degree to which these mechanisms explain perceptual changes, nor rule out additional mechanisms that could perform selection at a competitive stage.

The current results suggest that early modulations may be sufficient to account for the effects of temporal attention on perception. Importantly, they also argue against the idea that the behavioral effects of temporal attention result from better selection or maintenance of working memory representations (Shalev & van Ede, 2021), a possibility that is especially relevant for dynamic stimuli. Thus while visual discriminability is impaired by temporal competition, we find no evidence that attention regulates competition across time.

## Methods

### Participants

Eighteen participants (11 females, 7 males; mean age=26.4 years old, SD=5.3 years, based on self-report) completed the experiment. All participants, except authors J.A.M. and R.N.D., were naive to the study. The sample size was larger than those used in previous studies demonstrating effects of temporal attention (Denison et al., 2017, 2024). Based on a previous effect size (*η*_G_^2^=0.25; Denison et al., 2024), a power analysis using G*Power 3.1 (Faul et al., 2007) indicated that a sample size of 18 would achieve 96% power to detect an effect of attention on performance at a 0.05 significance level. No sex or gender-based analyses were performed, and we did not consider sex or gender in the study design, as neither sex nor gender played a role in our research questions. All participants had normal or corrected-to-normal vision and were monetarily compensated for their time. All participants provided informed consent and Boston University’s Institutional Review Board approved the experimental methods.

### Stimuli

Stimuli were generated using MATLAB R2023a and Psychophysics Toolbox 3.0.19 (Brainard, 1997; Kleiner et al., 2007; Pelli, 1997) on Linux. The stimuli were displayed on a VPixx VIEWPixx LCD display (VPixx Technologies Inc., QC, Canada) with a resolution of 1920 × 1080 and a refresh rate of 120 Hz and placed at a viewing distance of 75 cm. A chin-and-head rest stabilized the participants’ heads. Throughout the trial, a central white fixation circle with a diameter of 0.25° appeared on a gray background with mean luminance of 45.56 cd/m^2^. Stimulus timing and audiovisual synchrony were confirmed with photodiode and digital oscilloscope (PicoScope 2000 Series 2204A, Pico Technology, Cambridgeshire, UK) measurements to millisecond precision.

Visual stimuli were full contrast sinusoidal gratings with a Gaussian envelope (0.7° standard deviation), presented foveally for 50 ms each. Gratings had a spatial frequency of 4 cycles per degree, and one of four possible phases (0°, 90°, 180°, 270°). Gratings could be tilted clockwise or counterclockwise at participants’ individual tilt thresholds from either the vertical or horizontal axis. We varied the gratings in phase and axes to make it difficult to perform the orientation discrimination task by learning an image template and to discourage comparison between the sequential targets.

Auditory cues were pure sine wave tones, which could be high-pitched (1046.5 Hz, C6), low-pitched (440 Hz, A4), or the combination of the low and high tones. The pitch instructed participants to attend to a single time point (T1: high tone, T2: low tone) or to distribute attention across both time points (both tones). Auditory cues were 200 ms long, enveloped by cosine - squared ramps 10 ms in duration. Brief, auditory clicks (0.5 ms) generated as broadband noise ranging from 2–20 kHz were presented simultaneously with T1 and T2 onsets. Auditory stimuli were presented via Sennheiser HD 569 noise-isolating headphones (Sennheiser, Lower Saxony, Wedemark, DE) and a Focusrite Scarlett 2i2 audio interface (Focusrite, High Wycombe, Buckinghamshire, UK) at a comfortable volume.

### Task

Participants directed attention to different time points in a minimal stimulus sequence, while performing a challenging visual perceptual task **(Figure 2a)**. Each trial had two temporally predictable time points (T1 and T2) at which gratings could be presented. On each trial, gratings could be presented at both time points (creating temporal competition), at only one time point, or at neither time point. The probability of a grating’s occurrence at each time point was 50% and independent for each time point.

To manipulate voluntary temporal attention, a precue tone presented 1000 ms before T1 directed the participant’s attention to either T1 (high tone), T2 (low tone), or both time points (both tones played simultaneously). A response cue tone, presented 500 ms after T2, indicated a single time point to report. The precue matched the response cue on 60% of trials (valid trials), mismatched on 20% of trials (invalid trials), and was uninformative on 20% of trials (neutral trials), yielding 75% validity when a single time point was precued. Thus participants were incentivized to attend to the precued time point. T1 and T2 occurred at fixed, predictable times following the precue and were separated by a 250 ms stimulus onset asynchrony. To reduce temporal uncertainty about the possible target times, brief auditory clicks and fixation dims (from white to a light gray of 69.61 cd/m^2^, 0.5 ms) were presented simultaneously with T1 and T2, whether or not a grating appeared.

Participants reported whether or not they saw a grating at the response-cued time point and, if so, its tilt, resulting in three response options (grating clockwise, counterclockwise, or absent). Participants were allowed to make their response after a go cue, indicated by a fixation dim from white to a dark gray of 22.97 cd/m^2^, presented 600 ms after the response cue until the response was submitted. Participants had unlimited time to respond. When a response was registered, a fixation color change provided feedback (green=correct, red=incorrect) for 500 ms. After feedback, the screen was a blank gray except for the white fixation circle for 750 ms before the next trial. After each block of 40 trials, participants received block feedback in the form of percent accuracy.

Each participant completed 1280 trials across 32 blocks, over the course of two to three 1.5 hour sessions on separate days. Cue validity (valid, neutral, invalid), target presence (present, absent), non-target presence (present, absent), the response-cued time point (T1, T2), grating tilt (counterclockwise, clockwise), and grating axis (vertical, horizontal) were counterbalanced and independent. Grating phase was pseudorandomized, so that the probability of each phase’s occurrence was controlled but not its probability conditioned on other variables.

### Training

Before the main experiment, participants completed a training session to learn the task and to determine individual tilt thresholds. First, participants practiced trials identical to the main task, except that two gratings were always presented, each grating was tilted 5°, and all precues were neutral; participants repeated these trials until reaching 75% accuracy within a single block of 32 trials.

Then, to determine individual tilt thresholds, participants completed 128 trials identical to the main task, except precues were neutral, gratings were present or absent with equal probability, and tilt was adaptively adjusted trial-to-trial. Two staircases alternated, one starting from a coarse tilt (5°) and the other from a fine tilt (0.2°). Each staircase followed a 3-up-1-down procedure (converging to an accuracy of ∼79%); a 2-up-1-down procedure (converging to an accuracy of ∼71%) was used for three participants. Staircases were updated only if the target was present, as the tilt judgment was irrelevant when the target was absent. The threshold estimate from each staircase was the mean tilt over its last six trials, and the final threshold was taken as the mean of the two staircase estimates (mean=1.05°; SD=0.34°).

Participants then performed 64 practice trials using their individual tilt threshold, with gratings always present and all valid precues, followed by 64 trials identical to the main experiment.

### Online fixation monitoring

Fixation was monitored using an EyeLink 1000 Plus eye tracker (SR Research, Ottawa, Ontario, CA) with a sampling frequency of 1000 Hz. Trial initiation was contingent upon central fixation within a 2.5° radius for 300 ms. Fixation maintenance was required from the onset of the precue until the response cue. If fixation was broken during this time, the trial was stopped and repeated at the end of the block. On average, 3.43% (SD=2.97%) of trials were repeated per participant due to broken fixation.

### Signal detection theory analysis

We used signal detection theory (SDT) analysis (Green & Swets, 1966) to calculate sensitivity and criterion separately for detecting target presence from absence, and discriminating target orientation when present. For the detection measures, we considered a response of either counterclockwise or clockwise as reporting the target as “seen” because the alternative was to report the target as “absent”. Hits were defined as “seen” responses to a present target, and false alarms were defined as “seen” responses to an absent target. For the orientation discriminability measures, we used only trials in which the target was present and a tilt response was supplied because orientation information is not available otherwise. We arbitrarily defined “hits” as correctly identifying a clockwise tilted grating as clockwise, and false alarms as incorrectly reporting a counterclockwise tilted grating as clockwise.

To handle extreme hit or false alarm rates, for which the signal detection measures are indeterminate, we applied a 1/2N correction (Hautus et al., 2022), which adjusts rates of 0 to 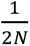 and rates of 1 to 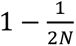, where *N* is the number of trials contributing to that rate.

Because the valid condition had three times more trials than the invalid and neutral conditions, differences in trial counts could affect SDT estimates, especially when accuracy is very high, as was the case in detecting full contrast gratings. To control trial counts across attention conditions, we performed a subsampling procedure as follows. For the detection measures, we randomly sampled trials without replacement from the valid condition per participant to match the trial counts for the invalid and neutral conditions. For the discrimination measures, we necessarily excluded trials in which an “absent” response was supplied, which could unequalize trial counts across SDT categories (i.e., the number of trials constituting “signal” versus “noise”). Therefore, for each participant, we identified the minimum number of trials within an SDT category across attention conditions, and subsampled all other categories to match this count. The subsampling procedure was repeated 1000 times for each measure and participant. The final trial-count-matched SDT measures were taken as the average of the estimates across subsamples.

### Statistical analysis

Repeated measures ANOVAs were used to assess the impact of experiment conditions on behavior using the *ez* package in R (Lawrence, 2016). For measures of accuracy, reaction time, and discrimination *d’* and criterion, the within-subject factors were validity (valid, neutral, or invalid with respect to the match between the precue and response cue), target time point (T1 or T2), target presence (present or absent), and non-target presence (present or absent). For measures of detection *d’* and criterion, the within-subject factors were validity, target time point, and non-target presence. When Mauchly’s test indicated a violation of sphericity assumptions (Mauchly, 1940), we reported Greenhouse-Geisser epsilon and corrected *p*-values. Effect sizes are provided as generalized eta squared.

When null results were of theoretical interest (i.e., in assessing the absence of an interaction between temporal attention and temporal competition), we quantified the amount of evidence in favor of the null versus alternative hypothesis using Bayes factor (BF_10_) (Lee & Wagenmakers, 2014). We calculated BF_10_ using the *BayesFactor* package in R (Morey & Rouder, 2023), by comparing a full model (H_1_), in which an interaction term of validity and non-target presence was included, against a reduced model (H_0_) that was identical except the interaction term was excluded. A BF_10_ greater than 1 indicates evidence in favor of the alternative whereas a BF_10_ less than 1 indicates evidence in favor of the null. To grade the decisiveness of evidence for the null hypothesis, we interpreted Bayes factors less than 1 following the recommendations of Lee and Wagenmakers (2014, Section 7.2).

## Declarations

## Availability of data and materials

The data are available at https://osf.io/kwfcx.

## Code availability

The code used to run the experiment and conduct the analyses presented here is available at https://github.com/denisonlab/ZOOT.

## Authors’ contributions

**K.J.T.:** Conceptualization, Methodology, Data Curation, Formal Analysis, Investigation, Visualization, Writing–Original Draft; **J.A.M.:** Conceptualization, Methodology, Data Curation, Formal Analysis, Investigation, Visualization, Writing–Original Draft; **R.N.D.:** Conceptualization, Supervision, Funding Acquisition, Project Administration, Writing– Reviewing & Editing.

