## Supplementary Material for "Temporal attention improves perception without enhancement by temporal competition"

Karen J. Tian*, Jennifer A. Motzer*, Rachel N. Denison
Department of Psychological and Brain Sciences, Boston University
*equal author contributions

For correspondence:


Department of Psychological and Brain Sciences
Boston University
111 Cummington Mall
Boston, MA 02215
United States

Keywords: temporal attention, temporal competition, selective attention, voluntary attention, visual perception

#### Supplementary Results

##### Temporal attentional effects on discrimination sensitivity

We assessed discrimination sensitivity for each target separately. Temporal attention enhanced discrimination sensitivity for each target when the non-target was present (T1: $F(2,34)=10.35$, $p<0.001$, $\eta_G^2=0.26$; T2: $F(2,34)=4.07$, $p=0.026$, $\eta_G^2=0.11$); however, when the non-target was absent, the attentional effect remained significant for T1 only (T1: $F(2,34)=4.09$, $p=0.026$, $\eta_G^2=0.08$; T2: $F(2,34)=1.00$, $p=0.379$, $\eta_G^2=0.01$, $BF_{10}=0.294$, indicating strong evidence in favor of the null).

One possible reason we did not observe a significant attentional effect on T2 discriminability without temporal competition was that T2 had higher performance overall (main effect of target: $F(1,17)=14.35$, $p<0.001$, $\eta_G^2=0.09$), leaving less opportunity for improvement with attention. To assess this possibility, we quantified each participant's attentional modulation as the difference in sensitivity between valid and invalid trials and correlated it to sensitivity on neutral trials, focusing on trials in which the non-target was absent. Across targets, discrimination performance on neutral trials was negatively correlated with the magnitude of attentional modulation across participants **(Figure S1)** ($r(16)=-0.458$, $p=0.005$). Assessing each target separately, this relationship remained significant for T2 ($r(16)=-0.539$, $p=0.021$) but not T1 ($r(16)=-0.325$, $p=0.188$). Future work using target-specific tilt titration procedures could compensate for such baseline performance differences across time.

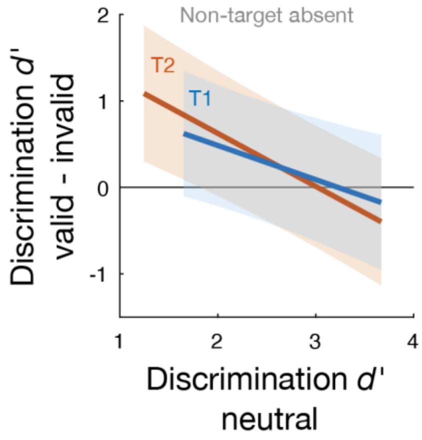


**Figure S1. Correlation between discrimination sensitivity on neutral trials and the magnitude of attentional benefits.** Across participants, discrimination sensitivity on neutral trials when the non-target was absent was negatively correlated with the magnitude of attentional modulation, suggesting that higher overall performance left less opportunity for improvement with attention. Data (n=18) are shown as the 95% confidence interval of the regression line.

### Supplementary Tables

#### **Supplementary Table 1.** Accuracy of the three-way perceptual judgment.

##### Target present

###### *Across targets*

| Effect | Target present |
|---|---|
| Target | $F(1,17)=69.12$, $p<0.001$, $\eta_G^2=0.23$ |
| Validity | $F(2,34)=28.74$, $p<0.001$, $\eta_G^2=0.15$ |
| NTP | $F(1,17)=0.08$, $p=0.787$, $\eta_G^2<0.01$ |
| Target × Validity | $F(2,34)=4.31$, $p=0.022$, $\eta_G^2=0.03$ |
| Target × NTP | $F(1,17)=0.81$, $p=0.381$, $\eta_G^2<0.01$ |
| Validity × NTP | $F(2,34)=0.06$, $p=0.941$, $\eta_G^2<0.01$ |
| Target × Validity × NTP | $F(2,34)=0.02$, $p=0.981$, $\eta_G^2<0.01$ |

###### *By target*

| Trials analyzed | Effect | T1 | T2 |
|---|---|---|---|
| *Across non-targets* | Validity | $F(2,34)=23.64$, $p<0.001$, $\eta_G^2=0.22$ | $F(2,34)=6.02$, $p=0.006$, $\eta_G^2=0.08$ |
| | NTP | $F(1,17)=0.04$, $p=0.848$, $\eta_G^2<0.01$ | $F(1,17)=1.01$, $p=0.328$, $\eta_G^2=0.01$ |
| | Validity × NTP | $F(2,34)=0.02$, $p=0.982$, $\eta_G^2<0.01$ | $F(2,34)=0.12$, $p=0.889$, $\eta_G^2<0.01$ |
| *Non-target present* | Validity | $F(2,34)=9.11$, $p<0.001$, $\eta_G^2=0.25$ | $F(2,34)=3.08$, $p=0.059$, $\eta_G^2=0.09$ |
| *Non-target absent* | Validity | $F(2,34)=11.42$, $p<0.001$, $\eta_G^2=0.19$ | $F(2,34)=5.04$, $p=0.012$, $\eta_G^2=0.07$ |

##### Target absent

###### *Across targets*

| Effect | Target absent |
|---|---|
| Target | $F(1,17)=3.03$, $p=0.100$, $\eta_G^2=0.01$ |
| Validity | $F(2,34)=2.33$, $p=0.113$, $\eta_G^2=0.01$ |
| NTP | $F(1,17)=21.31$, $p<0.001$, $\eta_G^2=0.30$ |
| Target × Validity | $F(2,34)=0.74$, $p=0.452$, $\eta_G^2<0.01$, $\varepsilon=0.76$ |
| Target × NTP | $F(1,17)=3.73$, $p=0.070$, $\eta_G^2=0.01$ |
| Validity × NTP | $F(2,34)=2.17$, $p=0.130$, $\eta_G^2<0.01$ |
| Target × Validity × NTP | $F(2,34)=1.06$, $p=0.358$, $\eta_G^2<0.01$ |

###### *By target*

| Trials analyzed | Effect | T1 | T2 |
|---|---|---|---|
| *Across non-targets* | Validity | $F(2,34)=2.61$, $p=0.088$, $\eta_G^2=0.01$ | $F(2,34)=0.68$, $p=0.512$, $\eta_G^2<0.01$ |
| | NTP | $F(1,17)=13.84$, $p=0.002$, $\eta_G^2=0.25$ | $F(1,17)=23.72$, $p<0.001$, $\eta_G^2=0.34$ |
| | Validity × NTP | $F(2,34)=1.74$, $p=0.190$, $\eta_G^2=0.01$ | $F(2,34)=1.26$, $p=0.292$, $\eta_G^2=0.01$, $\varepsilon=0.75$ |
| *Non-target present* | Validity | $F(2,34)=2.30$, $p=0.116$, $\eta_G^2=0.02$ | $F(2,34)=0.97$, $p=0.389$, $\eta_G^2=0.01$ |
| *Non-target absent* | Validity | $F(2,34)=0.51$, $p=0.537$, $\eta_G^2=0.02$, $\varepsilon=0.68$ | $F(2,34)=1.24$, $p=0.289$, $\eta_G^2=0.04$, $\varepsilon=0.62$ |

Factors of target (T1, T2) × validity (valid, neutral, invalid) × non-target presence (NTP; present, absent) in the full design. Separate ANOVAs include only factors that varied within the indicated subsets. Degrees of freedom are uncorrected; where Mauchly’s test indicated a violation of sphericity, $p$ values are Greenhouse–Geisser corrected and $\varepsilon$ is reported.

#### **Supplementary Table 2.** Reaction times.

#### Target present

##### *Across targets*

| Effect | Target present |
|---|---|
| Target | $F(1,17)=24.53$, $p<0.001$, $\eta_G^2=0.03$ |
| Validity | $F(2,34)=18.20$, $p<0.001$, $\eta_G^2=0.07$, $\varepsilon=0.67$ |
| NTP | $F(1,17)=0.32$, $p=0.578$, $\eta_G^2<0.01$ |
| Target × Validity | $F(2,34)=5.15$, $p=0.011$, $\eta_G^2<0.01$ |
| Target × NTP | $F(1,17)=1.75$, $p=0.203$, $\eta_G^2<0.01$ |
| Validity × NTP | $F(2,34)=0.14$, $p=0.867$, $\eta_G^2<0.01$ |
| Target × Validity × NTP | $F(2,34)=1.01$, $p=0.375$, $\eta_G^2<0.01$ |

##### *By target*

| Trials analyzed | Effect | T1 | T2 |
|---|---|---|---|
| | Validity | $F(2,34)=16.48$, $p<0.001$, $\eta_G^2=0.09$ | $F(2,34)=15.04$, $p<0.001$, $\eta_G^2=0.06$, $\varepsilon=0.59$ |
| *Across non-targets* | NTP | $F(1,17)=0.16$, $p=0.696$, $\eta_G^2<0.01$ | $F(1,17)=1.57$, $p=0.228$, $\eta_G^2<0.01$ |
| | Validity × NTP | $F(2,34)=0.22$, $p=0.802$, $\eta_G^2<0.01$ | $F(2,34)=0.71$, $p=0.500$, $\eta_G^2<0.01$ |
| *Non-target present* | Validity | $F(2,34)=16.83$, $p<0.001$, $\eta_G^2=0.10$ | $F(2,34)=9.91$, $p=0.004$, $\eta_G^2=0.07$, $\varepsilon=0.59$ |
| *Non-target absent* | Validity | $F(2,34)=11.57$, $p<0.001$, $\eta_G^2=0.09$ | $F(2,34)=12.98$, $p<0.001$, $\eta_G^2=0.04$ |

#### Target absent

##### *Across targets*

| Effect | Target absent |
|---|---|
| Target | $F(1,17)=12.44$, $p=0.003$, $\eta_G^2=0.02$ |
| Validity | $F(2,34)=5.42$, $p=0.009$, $\eta_G^2=0.02$ |
| NTP | $F(1,17)=35.70$, $p<0.001$, $\eta_G^2=0.14$ |
| Target × Validity | $F(2,34)=1.45$, $p=0.248$, $\eta_G^2<0.01$ |
| Target × NTP | $F(1,17)=5.96$, $p=0.026$, $\eta_G^2=0.01$ |
| Validity × NTP | $F(2,34)=10.62$, $p<0.001$, $\eta_G^2=0.02$ |
| Target × Validity × NTP | $F(2,34)=1.28$, $p=0.285$, $\eta_G^2<0.01$, $\varepsilon=0.70$ |

##### *By target*

| Trials analyzed | Effect | T1 | T2 |
|---|---|---|---|
| | Validity | $F(2,34)=2.82$, $p=0.074$, $\eta_G^2=0.01$ | $F(2,34)=5.53$, $p=0.008$, $\eta_G^2=0.03$ |
| *Across non-targets* | NTP | $F(1,17)=25.81$, $p<0.001$, $\eta_G^2=0.18$ | $F(1,17)=31.90$, $p<0.001$, $\eta_G^2=0.09$ |
| | Validity × NTP | $F(2,34)=4.86$, $p=0.014$, $\eta_G^2=0.01$ | $F(2,34)=8.47$, $p=0.004$, $\eta_G^2=0.03$, $\varepsilon=0.72$ |
| *Non-target present* | Validity | $F(2,34)=5.88$, $p=0.006$, $\eta_G^2=0.03$ | $F(2,34)=15.14$, $p<0.001$, $\eta_G^2=0.09$ |
| *Non-target absent* | Validity | $F(2,34)=0.07$, $p=0.930$, $\eta_G^2<0.01$ | $F(2,34)=0.04$, $p=0.915$, $\eta_G^2<0.01$, $\varepsilon=0.72$ |

Factors of target (T1, T2) × validity (valid, neutral, invalid) × non-target presence (NTP; present, absent) in the full design. Separate ANOVAs include only factors that vary within the indicated subsets. Degrees of freedom are uncorrected; where Mauchly's test indicated a violation of sphericity, *p* values are Greenhouse-Geisser corrected and ε is reported.

**Supplementary Table 3.** Discrimination sensitivity ($d'$).

***Discrimination sensitivity (across targets)***

| Effect | Discrimination $d'$ |
|---|---|
| Target | $F(1,17)=49.36$, $p<0.001$, $\eta_G^2=0.14$ |
| Validity | $F(2,34)=19.44$, $p<0.001$, $\eta_G^2=0.10$ |
| NTP | $F(1,17)=22.61$, $p<0.001$, $\eta_G^2=0.13$ |
| Target × Validity | $F(2,34)=1.59$, $p=0.219$, $\eta_G^2=0.01$ |
| Target × NTP | $F(1,17)=3.37$, $p=0.084$, $\eta_G^2=0.01$ |
| Validity × NTP | $F(2,34)=2.45$, $p=0.101$, $\eta_G^2=0.02$ |
| Target × Validity × NTP | $F(2,34)=0.27$, $p=0.764$, $\eta_G^2<0.01$ |

***Discrimination sensitivity (by target)***

| Trials analyzed | Effect | T1 | T2 |
|---|---|---|---|
| | Validity | $F(2,34)=17.22$, $p<0.001$, $\eta_G^2=0.17$ | $F(2,34)=4.82$, $p=0.014$, $\eta_G^2=0.05$ |
| *Across non-targets* | NTP | $F(1,17)=20.92$, $p<0.001$, $\eta_G^2=0.20$ | $F(1,17)=10.32$, $p=0.005$, $\eta_G^2=0.07$ |
| | Validity × NTP | $F(2,34)=1.70$, $p=0.197$, $\eta_G^2=0.03$ | $F(2,34)=1.53$, $p=0.231$, $\eta_G^2=0.02$ |
| *Non-target present* | Validity | $F(2,34)=10.35$, $p<0.001$, $\eta_G^2=0.26$ | $F(2,34)=4.07$, $p=0.026$, $\eta_G^2=0.11$ |
| *Non-target absent* | Validity | $F(2,34)=4.09$, $p=0.026$, $\eta_G^2=0.08$ | $F(2,34)=1.00$, $p=0.379$, $\eta_G^2=0.01$ |

Factors of target (T1, T2) × validity (valid, neutral, invalid) × non-target presence (NTP; present, absent) in the full design. Separate ANOVAs include only factors that vary within the indicated subsets. Degrees of freedom are uncorrected; where Mauchly's test indicated a violation of sphericity, *p* values are Greenhouse-Geisser corrected and ε is reported.

**Supplementary Table 4.** Detection sensitivity (*d′*) and criterion.

##### *Detection sensitivity (across targets)*

| Effect | Detection *d′* |
|---|---|
| Target | $F(1,17)=5.78$, $p=0.028$, $\eta_G^2=0.02$ |
| Validity | $F(2,34)=1.96$, $p=0.157$, $\eta_G^2=0.01$ |
| NTP | $F(1,17)=6.12$, $p=0.024$, $\eta_G^2=0.05$ |
| Target × Validity | $F(2,34)=1.31$, $p=0.282$, $\eta_G^2<0.01$ |
| Target × NTP | $F(1,17)=16.50$, $p<0.001$, $\eta_G^2=0.09$ |
| Validity × NTP | $F(2,34)=11.10$, $p=0.001$, $\eta_G^2=0.03$, $\varepsilon=0.71$ |
| Target × Validity × NTP | $F(2,34)=1.96$, $p=0.156$, $\eta_G^2=0.01$ |

##### *Detection sensitivity (by target)*

| Trials analyzed | Effect | T1 | T2 |
|---|---|---|---|
| *Across non-targets* | Validity | $F(2,34)=3.26$, $p=0.051$, $\eta_G^2=0.02$ | $F(2,34)=0.64$, $p=0.534$, $\eta_G^2=0.01$ |
| | NTP | $F(1,17)=0.38$, $p=0.544$, $\eta_G^2<0.01$ | $F(1,17)=17.29$, $p<0.001$, $\eta_G^2=0.24$ |
| | Validity × NTP | $F(2,34)=6.64$, $p=0.008$, $\eta_G^2=0.06$, $\varepsilon=0.75$ | $F(2,34)=2.49$, $p=0.098$, $\eta_G^2=0.02$ |
| *Non-target present* | Validity | $F(2,34)=1.60$, $p=0.217$, $\eta_G^2=0.02$ | $F(2,34)=0.18$, $p=0.835$, $\eta_G^2<0.01$ |
| *Non-target absent* | Validity | $F(2,34)=9.06$, $p=0.003$, $\eta_G^2=0.15$, $\varepsilon=0.72$ | $F(2,34)=5.86$, $p=0.006$, $\eta_G^2=0.11$ |

##### Detection criterion *(across targets)*

| Effect | Detection criterion |
|---|---|
| Target | $F(1,17)=19.34$, $p<0.001$, $\eta_G^2=0.10$ |
| Validity | $F(2,34)=12.52$, $p<0.001$, $\eta_G^2=0.04$ |
| NTP | $F(1,17)=66.39$, $p<0.001$, $\eta_G^2=0.58$ |
| Target × Validity | $F(2,34)=2.56$, $p=0.092$, $\eta_G^2=0.01$ |
| Target × NTP | $F(1,17)=4.54$, $p=0.048$, $\eta_G^2=0.01$ |
| Validity × NTP | $F(2,34)=2.18$, $p=0.129$, $\eta_G^2=0.01$ |
| Target × Validity × NTP | $F(2,34)=0.40$, $p=0.671$, $\eta_G^2<0.01$ |

##### Detection criterion (*by target)*

| Trials analyzed | Effect | T1 | T2 |
|---|---|---|---|
| *Across non-targets* | Validity | $F(2,34)=9.75$, $p<0.001$, $\eta_G^2=0.08$ | $F(2,34)=1.25$, $p=0.300$, $\eta_G^2=0.01$ |
| | NTP | $F(1,17)=57.87$, $p<0.001$, $\eta_G^2=0.61$ | $F(1,17)=58.15$, $p<0.001$, $\eta_G^2=0.55$ |
| | Validity × NTP | $F(2,34)=1.19$, $p=0.317$, $\eta_G^2=0.01$ | $F(2,34)=1.44$, $p=0.252$, $\eta_G^2=0.02$ |
| *Non-target present* | Validity | $F(2,34)=2.54$, $p=0.093$, $\eta_G^2=0.04$ | $F(2,34)=0.13$, $p=0.876$, $\eta_G^2<0.01$ |
| *Non-target absent* | Validity | $F(2,34)=8.42$, $p=0.001$, $\eta_G^2=0.17$ | $F(2,34)=4.08$, $p=0.026$, $\eta_G^2=0.12$ |

Factors of target (T1, T2) × validity (valid, neutral, invalid) × non-target presence (NTP; present, absent) in the full design. Separate ANOVAs include only factors that vary within the indicated subsets. Degrees of freedom are uncorrected; where Mauchly's test indicates a violation of sphericity, *p* values are Greenhouse-Geisser corrected and ε is reported.
